# Ldb1 mediates *trans* enhancement in mammals

**DOI:** 10.1101/287524

**Authors:** K. Monahan, A. Horta, A.M. Mumbay-Wafula, L. Li, Y. Zhao, P.E. Love, S. Lomvardas

## Abstract

Singular olfactory receptor (OR) gene expression^1,2^ coincides with the formation of a multi-chromosomal enhancer hub that associates with the only transcribed OR allele in each cell^3,4^. This hub consists of converging transcriptional enhancers^3^, or “Greek Islands”, defined by stereotypic binding of Lhx2 and Ebf on a shared, composite DNA motif^5^. How this multi-chromosomal hub, or any other genomic compartment, assembles is unknown, and so is the significance of compartmentalization in transcription. Here, we report that LIM domain binding protein 1 (Ldb1), which is recruited by Lhx2 and Ebf to Greek Islands, promotes robust and specific *trans* interactions between these enhancers. In addition to disrupting Greek Island hubs, Ldb1 deletion also causes significant downregulation of OR transcription. Thus, our data provide insight to the formation of genomic compartments, confirm the essential role of interchromosomal interactions in OR gene choice, and establish *trans* enhancement as a mechanism for mammalian gene activation.

---

To obtain insight into the formation of Greek Island hubs we examined the genomic distribution of two known regulators of genomic organization, Ctcf and Rad21^6–9^. ChIP-seq in mature olfactory sensory neurons (mOSNs), revealed very little binding of either protein to Greek Islands (Fig.1a, b). Thus, we explored other mediators of genomic interactions, namely the Ldb family proteins^10–14^, which interact with LIM-domain proteins, such as Lhx2^15^. Ldb1, which is the only Ldb family member expressed in OSNs (Extended data Fig.1a, b), has ~22,000 ChIP-seq peaks in mOSNs that closely overlap with Lhx2 peaks (Extended data Fig. 2). Consistent with this genomewide pattern, every one of the 63 Lhx2-bound Greek Islands coincide with an Ldb1 peak (Fig.1a, b). Ldb1 ChIP-seq from Lhx2 KO mOSNs shows that Ldb1 recruitment to the Greek Islands, and to most of its other targets, is Lhx2 dependent (Fig.1c). However, because Lhx2 deletion also reduces Ebf binding to the Greek Islands^5^, we cannot rule out a synergistic interaction between Lhx2 and Ebf in Ldb1 recruitment. This could explain why Greek Islands represent some of the strongest Ldb1 peaks in mOSNs (Fig.1b, d), and why Lhx2/Ebf co-bound peaks are stronger than Lhx2-only peaks genomewide (Fig.1e). Moreover, Ldb1 ChIP-seq peaks that are preserved, or gained, following Lhx2 deletion, are frequently Ebf bound and have Ebf DNA binding motif as the most enriched consensus motif (Extended data Fig.3).

**Figure 1:**
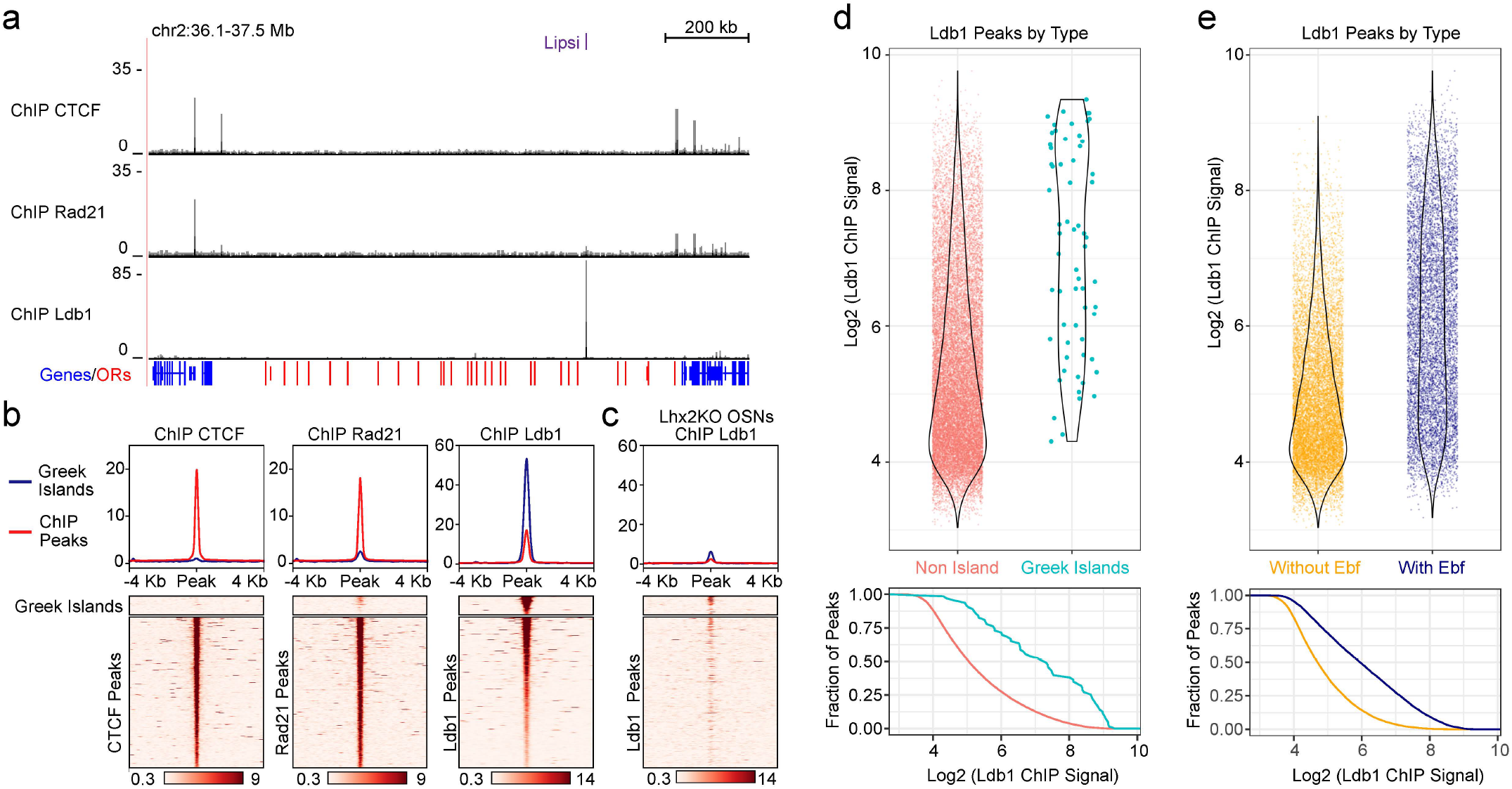
Ldb1 binds Greek Islands. **a)** OSN ChIP-seq signal for CTCF, Rad21, and Ldb1 across the OR gene cluster containing the Greek Island Lipsi. OR genes are red and all other genes are blue. **b)** mOSN ChIP signal over Greek Islands and genomewide ChIP peaks. Plot shows average signal across all sites and heatmap shows individual peaks. **c)** Ldb1 ChIP signal from Lhx2KO mOSNs. **d)** Upper: Normalized ChIP signal per peak for Greek Islands versus all other Ldb1 peaks. Lower: fraction of peaks with signal greater than value on x. **e)** as in D, except grouping Ldb1 peaks by overlap with Ebf ChIP peaks.

To explore the function of Ldb1 in Greek Island interactions we deleted Lbd1 in mOSNs, by crossing a conditional Ldb1 allele^16^ to Omp-ires-Cre mice^17^ (Fig. 2a). This genetic strategy eliminated Ldb1 from the more apical, mOSN layer of the MOE, but not from less differentiated OSNs and progenitor cells (Fig. 2b). To allow purification of Ldb1 KO mOSNs in HiC experiments, we included a Cre-inducible fluorescent reporter (TdTomato)^18^ in our genetic crosses (Fig.2a). *In situ* HiC on control and Ldb1 KO FAC-sorted mOSNs did not reveal quantitative changes in genomic organization, with genomewide interchromosomal contacts being almost as frequent as in control mOSNs (35.6% interchromosomal contacts in control mOSNs vs 32.8% interchromosomal contacts in Ldb1 KO mOSNs, Extended Data Fig. 4a,b). Consistently, OR gene clusters continue to contact each other with high specificity (Fig. 2c), and HMM-based compartment prediction^19^ identifies the same OR-enriched compartment in Ldb1 KO and control mOSNs (Fig. 2d). Finally, at the primary level of genomic organization, the topologically associated domains (TADs)^20,21^, we observe no effect of Ldb1 deletion on the partitioning of OR gene clusters (Fig. 2d, Extended Data Fig.4b, 5c).

**Figure 2:**
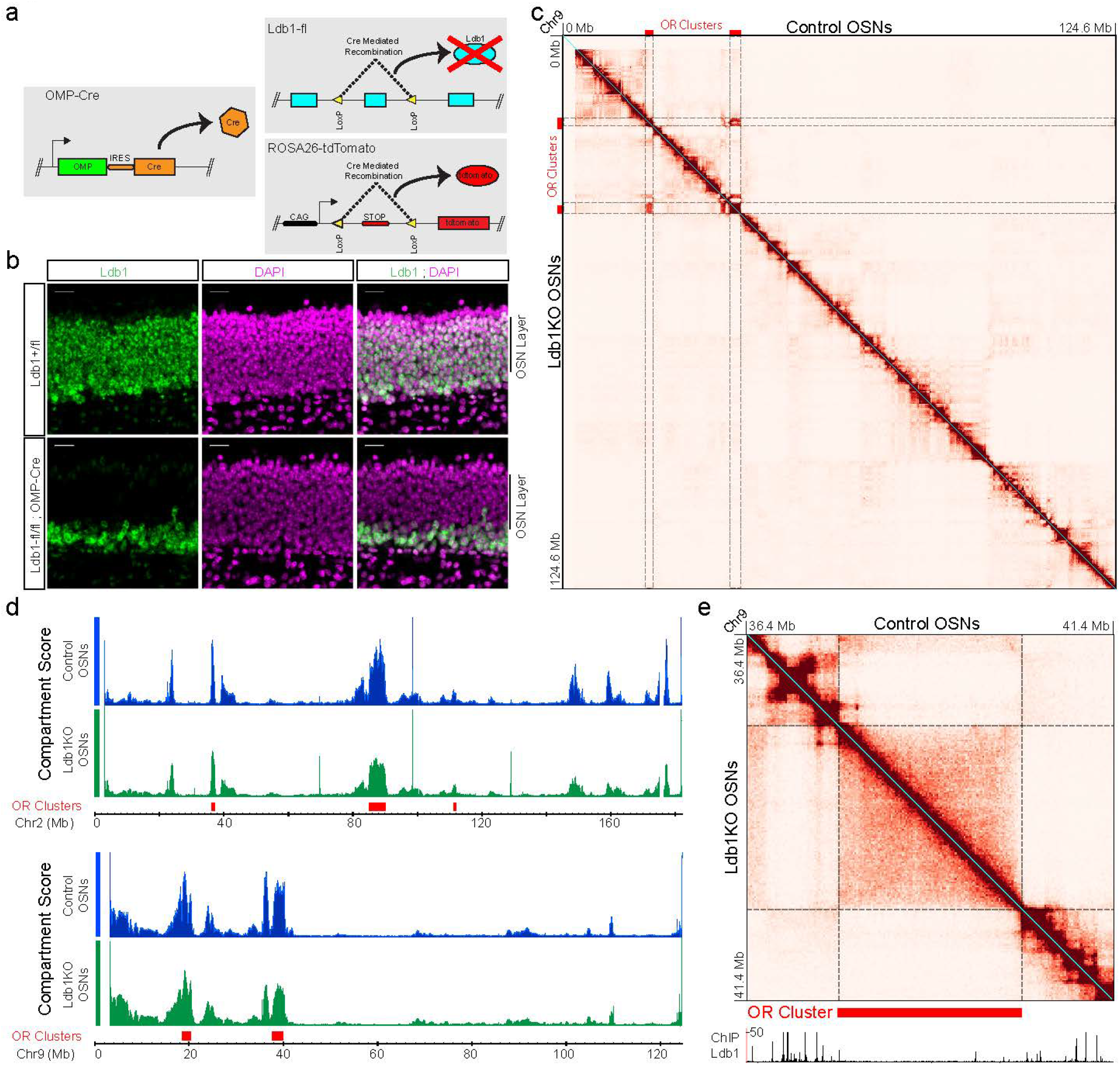
Ldb1KO does not affect overall nuclear architecture. **a)** Genetic strategy for conditional deletion and fluorescent labeling. **b)** In Ldb1fl/fl;OMP-Cre mice, Ldb1 (green) is lost from mature OSNs but retained in basal immature cells. Nuclei are stained with DAPI (magenta). Scale bar = 20um **c)** HiC profile of chromosome 9 is similar for Control and Ldb1KO OSNs (100kb resolution, max=250 contacts per billion). **d)** Compartment analysis identifies a similar OR cluster associated compartment in Control and Ldb1KO OSNs. **e)** Local pattern of HiC contacts around OR clusters on chromosome 9 is retained in Ldb1KO OSNs (25kb resolution, max = 100 contacts per billion). Ldb1 ChIP signal from control OSNs is shown below the contact map.

In sharp contrast to the weak effects of Ldb1 deletion on the topology and compartmentalization of OR clusters, genomic interactions between Greek Islands are drastically reduced. Visual examination of *trans* interactions between OR clusters shows that HiC interaction “hotspots” centered on Greek Islands disappear upon Ldb1 deletion (Fig. 3a). The reduction of interactions between Greek Islands is widespread, as pairwise or aggregate interaction frequencies between the 63 Greek Islands show strong reduction of these contacts throughout the Greek Island repertoire (Fig. 3b,c Extended data Fig.5, 6a). Aggregate peak analysis (APA)^19^, also shows that Ldb1 deletion disrupts the highly focal *trans* contacts between Greek Islands (Fig. 3d). Importantly, it should be noted that Ldb1 deletion does not prevent Greek Islands from making genomic contacts altogether; instead it reduces their contact frequencies to the level of surrounding OR sequences (Fig. 3e, extended data Fig.5a, b). We do note, however, a reduction in long distance contacts between OR clusters, but the magnitude of this reduction is much smaller than for contacts between Greek Islands (Fig, 3e, Extended Data Fig.5c, d).

**Figure 3:**
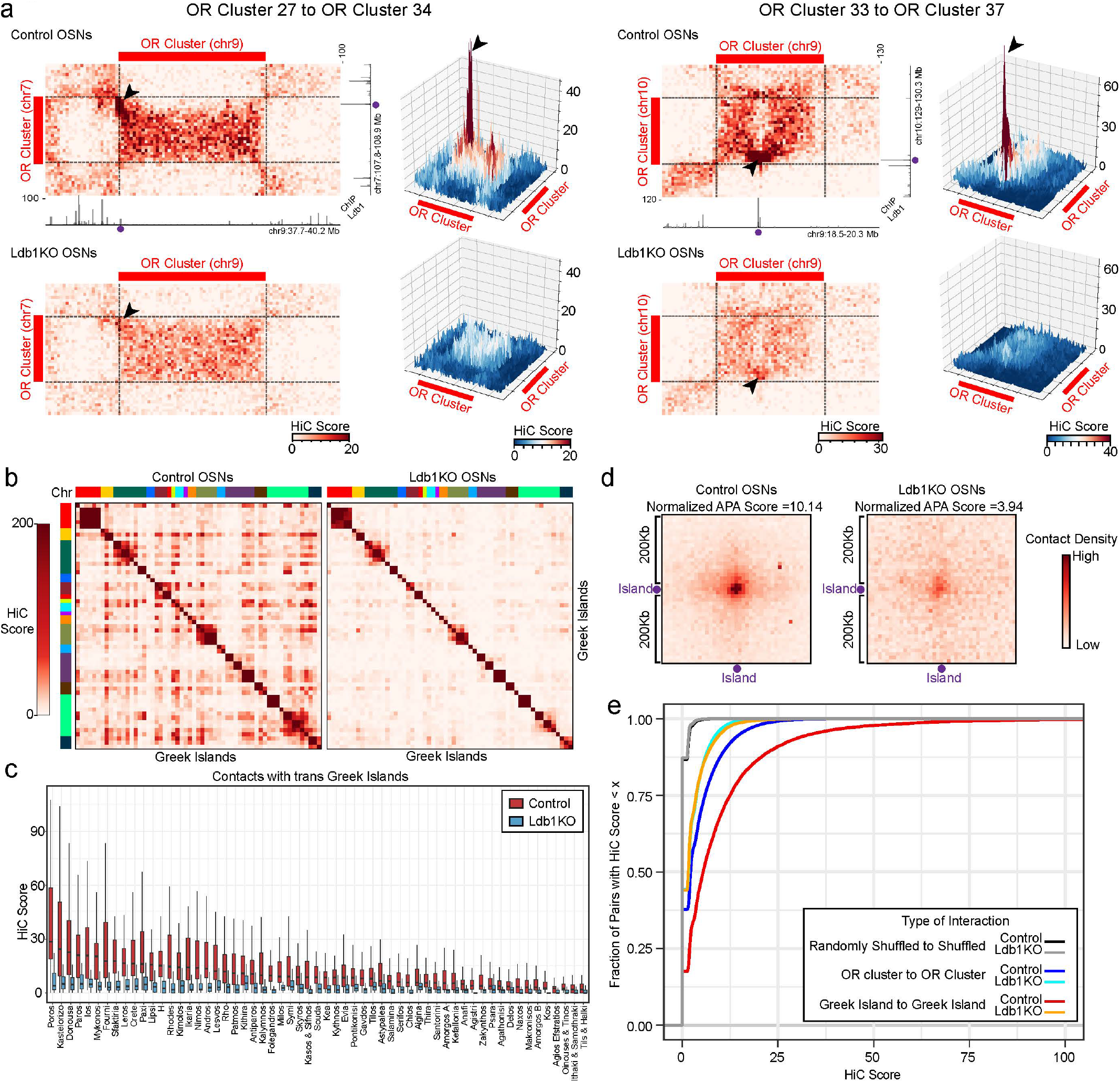
Deletion of Ldb1 results in dissolution of long range contacts between Greek Islands. **a)** HiC contacts between pairs of OR clusters located on different chromosomes (50kb resolution). Ldb1 ChIP-seq signal is shown for control OSNs. Interacting Greek Islands are marked with purple dots and their contact points are indicated by arrowheads on the contact matrix. **b)** Pairwise HiC contacts between all pairs of Greek Islands ordered by genomic position in Control (left) and Ldb1 KO (right) mOSNs **c)** Distribution of contacts made by each Greek Island with all other Greek Islands located on different chromosomes **d)** Aggregate peak analysis (APA) of *trans* Greek Island contacts. Degree of central enrichment indicates strength of interactions. **e)** Effect of Ldb1KO on distribution of HiC scores for different classes of *trans* interactions.

To examine the role of Ldb1-mediated genomic interactions in OR transcription, we first focused our analysis on Olfr1507, which is the most proximal OR gene to the Greek Island H^22^. Although transcription of this OR depends on H^23,24^, our accompanying paper^4^ and previous studies^3^ show that the transcriptionally active Olfr1507 also makes extensive contacts with a large number of Greek Islands from other chromosomes. Importantly, Ldb1 deletion has little effect on local contacts between H and the Olfr1507 cluster, while it eliminates *trans* interactions between H and the other Greek Islands (Extended Data Fig. 6a,b), providing an ideal system for dissecting the contribution of *trans* interactions in transcription. In support of the essential function of interchromosomal contacts in Olfr1507 expression, Olfr1507 protein is only detected in basal, Ldb1-positive cells in sections of Ldb1 KO MOEs, while it is absent from apical mOSNs that deleted Ldb1 (Fig. 4a). RNA-seq analysis demonstrates that the presumed reduction of Olfr1507 transcript levels in Ldb1 KO mOSNs is strong and highly significant (18.48-fold reduction, p_adj_=9.34e-47) (Fig.4b). Furthermore, RNA-seq shows that the significant downregulation of OR transcription extends to most ORs in the genome with 67% of OR genes significantly downregulated with p_adj_<0.01 (Fig. 4c). The few ORs that are not downregulated by Ldb1 deletion are predominantly type I, “fish-like” ORs (extended data Fig. 6c), which are evolutionarily older, lack heterochromatic marks, and are subject to different regulatory mechanisms^25–27^. Finally, we find that Ldb1 deletion has a relatively limited impact on the expression of non-OR genes (Fig.4c). Even when we focus our analysis on the genes with Ldb1 peaks on their promoter or proximal enhancers, most genes are not significantly affected by Ldb1 deletion and these genes are only slightly more likely to be significantly upregulated or downregulated than genes that are not associated with Ldb1 (Fig.4d). Thus, although Ldb1 has diverse and essential functions during development, in post-mitotic mOSNs its requirement in transcription strongly correlates with its role as mediator of *trans* enhancer interactions.

**Figure 4:**
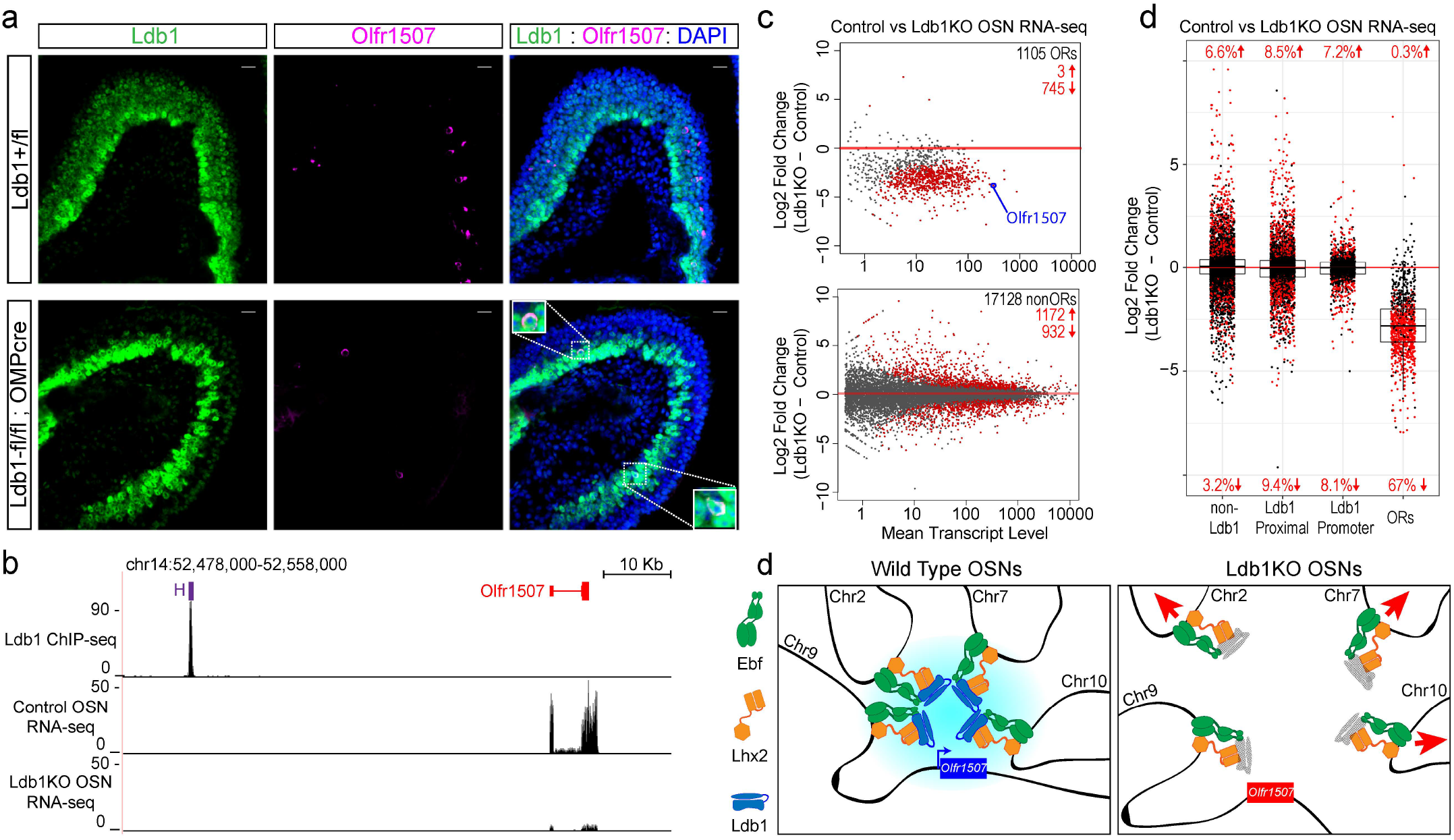
Loss of Ldb1 reduces OR expression. **a)** Olfr1507 (magenta) positive cells in Ldb1KO mice are restricted to basal immature OSNs that are still Ldb1 (green) immunoreactive. Nuclei are stained with DAPI (blue). Scale bar = 20um. **b)** Greek Island H is strongly bound by Ldb1, and loss of Ldb1 results in reduced levels of Olfr1507 by RNA-seq. **c)** RNA-seq analysis of gene expression in Ldb1KO OSNs relative to Control OSNs. Significantly changed genes (p_adj_ < 0.01) are in red **d)** Effect of Ldb1KO on genes not associated with Ldb1 ChIP peaks, genes located closest to a non-promoter Ldb1 ChIP-seq peak, genes with an Ldb1 ChIP-seq peak within the promoter region, and ORs. **e)** Lhx2 and Ebf recruit Ldb1 to Greek Islands. Ldb1 drives the formation of long range and interchromosomal interactions with other Greek Islands. Assembly of Greek Islands into a hub activates expression of an OR. Deletion of Ldb1 results in dissolution of the Greek Island hub and loss of OR expression.

Here, we identify Ldb1 as the first protein involved in *trans* genomic interactions and compartmental segregation, and reveal the essential role of interchromosomal compartments in mammalian gene expression. Ldb1 is recruited to the Greek Islands by Lhx2 and, likely, Ebf and regulates their extensive interchromosomal interactions, assembling a multi-enhancer hub that appears essential for OR transcription (Fig. 4e). Given the common genetic and biochemical valance of the Greek Islands, this 3-dimensional super-enhancer could be viewed as a “polymer” that deploys the synergistic action of multiple weak “monomers”, the individual Greek Islands. Consistent with the recently proposed role of phase transitions in compartmental segregation^28^ and super-enhancer function^29^, this model could explain why OR activation requires such an extraordinary number of interacting enhancers: Since phase transitions occurs in concentration dependent fashion^30^, a distinct OR-activating phase may form only upon accumulation of sufficient Lhx2/Ebf/Ldb1 complexes, delivered locally by converging Greek Islands. If individual Greek Islands cannot activate OR transcription on their own, then only the single OR allele that associates with the Greek Island hub (see accompanying paper^4^) will be transcriptionally active, providing a solution to the absolute singularity of OR transcription. Moreover, because this multi-enhancer hub is not pre-assembled at any given chromosomal location, like canonical super-enhancers, distribution of its individual components into 18 different chromosomes prevents transcriptional domination of the MOE by a “privileged”, hub-proximal OR. Thus, the dispersion of Greek Islands into multiple chromosomes, their inability to activate OR transcription on their own, and their tendency to interact in *trans*, satisfies key requirements for a sensitive, widely tuned, and evolutionarily plastic olfactory system: robust transcriptional activation of the chosen OR; low, distributed frequency of transcriptional activation for any particular OR allele; and biologically irrelevant transcriptional rates for non-chosen ORs. This novel regulatory paradigm of Ldb-mediated *trans* enhancement may serve as generator of molecular diversity in any biological system seeking transcriptional diversity through stochastic, mutually exclusive choices.

## Acknowledgments

We would like to thank Ira Schieren for flow cytometry and members of the Lomvardas lab for input, suggestions, and discussions and for critical reading of the manuscript. KM was funded by F32 post-doctoral fellowship GM108474 (NIH), and AH was funded by F31 post-doctoral fellowship DC016785 (NIH). This project was funded by U01DA0408052, R01DC013560 and R01DC015451 (NIH), and the HHMI Faculty Scholar Award. Research reported in this publication was performed in the CCTI Flow Cytometry Core, supported in part by the Office of the Director, National Institutes of Health under awards S10RR027050. The content is solely the responsibility of the authors and does not necessarily represent the official views of the National Institutes of Health.

## Materials and Methods

### Animal Models

Mice were treated in compliance with the rules and regulations of IACUC under protocol number AC-AAAT2450. Mice were sacrificed using CO_2_ followed by cervical dislocation. All experiments were performed on dissected olfactory epithelium tissue or on dissociated cells prepared from whole olfactory epithelium tissue. Dissociated cells were prepared using papain (Worthington Biochemical) and FAC sorted as previously described^5^.

Control mature olfactory sensory neurons (mOSNs) were FACS purified from *OMP-ires-GFP* mice^31^. Conditional deletion of Ldb1 in OSNs was achieved by crossing Ldb1 conditional allele mice (Ldb1-fl: Ldb1^tm2Lmgd^)^16^ with *OMP-ires-Cre* mice (OMP-Cre, Omp^tm1(cre)Jae^)^17^. For experiments on purified Ldb1KO mOSNs, a Cre-inducible tdtomato allele (ROSA26-tdtomato, Gt(ROSA)26Sor^tm14(CAG-tdTomato)Hze/J^)^18^ was included in the cross and tdtomato+ cells were purified by FACS as previously described^5^. Conditional deletion of Lhx2 in mOSNs was achieved by crossing *Lhx2* conditional allele mice (Lhx2-fl: Lhx2^tm1Monu^)^32^ with OMP-Cre mice. Lhx2KO mOSNs were purified by including ROSA26-tdtomato and purifying tdtomato+ cells as previously described^5^.

### Chromatin Immunoprecipitation

Chromatin Immunoprecipitation (ChIP) experiments were carried out as previously described^5^. Briefly, dissociated cells were fixed for 5 minutes with 1% formaldehyde in PBS and then FAC sorted to purify OSNs. Sheared chromatin was prepared from FACS purified cells using a Covaris S220 Focused-ultrasonicator. ChIP was performed using antibodies for CTCF (Millipore Cat# 07-729, RRID:AB_441965), Rad21 (Abcam Cat# ab992, RRID:AB_2176601), or Ldb1 (Santa Cruz Biotechnology Cat# sc-11198, RRID:AB_2137017). ChIP-seq libraries were prepared using the Nugen Ovation Ultralow Library System v2 (Nugen Cat# 0344-32). See ChIP-seq tab of Supporting Information 1 for a summary of ChIP-seq data sets.

Adapter sequences were removed from raw ChIP-seq data using CutAdapt^33^ (RRID:SCR_011841) and filtered reads were aligned to the mouse genome (mm10) using Bowtie2^34^ v2.3.2 (RRID:SCR_006646) with default settings. Picard (RRID:SCR_006525) was used to identify duplicate reads, which were then removed with Samtools^35^ (RRID:SCR_002105) (Li et al., 2009). Samtools was used to select uniquely aligning reads by removing reads with alignment quality alignments below 30 (-q 30). Peaks of ChIP-seq signal were identified using HOMER^36^ (RRID:SCR_010881) in “factor” mode with an input control. Consensus peak sets were generated by selecting peaks that overlapped in at least two biological replicates and extending them to their combined size. Bedtools2^37^ was used to compare peak sets (Quinlan and Hall, 2010). Motif discovery and analysis was performed with HOMER.

For signal tracks, biological replicates were merged and HOMER was used to generate 1bp resolution signal tracks normalized to a library size of 10,000,000 reads. Values in all ChIP-seq signal plots are reads per 10 million. Plots of ChIP-seq signal over individual loci were generated using the UCSC Genome Browser. Deeptools2^38^ was used to generate ChIP-seq heatmaps and mean signal plots. Each row of the heatmap is an 8kb region centered on a Greek Island or ChIP-seq peak for the factor shown. For heatmap in Figure 1b and 1c, all Greek Islands are shown alongside 500 randomly selected ChIP-seq peaks for each factor. Signal plots present average data for all peaks. Heatmaps are sorted by mean signal, except for extended-data figure 3 where all heatmaps are sorted by Ldb1 ChIP-seq signal in Control OSNs.

DiffBind^39^ was used to calculate ChIP-seq signal in each peak. For this analysis, Diffbind was used to normalize ChIP-seq scores across biological replicate experiments using the “DBA_SCORE_TMM_READS_EFFECTIVE” scoring system, which normalizes using edgeR and the effective library size. The ChIP-seq signal for each peak was then calculated by averaging the normalized score across biological replicates.

### HiC

HiC experiments were performed and sequencing data aligned as described in Horta et al^4^. See Supporting Information 2 for a summary of HiC data sets.

HiC figures were prepared with Juicebox v1.8.8 or using HiC scores extracted with Juicer Tools^40^. HiC score shown are KR normalized counts scaled to a library size of 1 billion HiC contacts (contacts per billion). Unless otherwise noted, plots show data at 50kb resolution. Greek Islands and OR clusters on the X chromosome were excluded from HiC analysis to avoid effects of X-inactivation. Compartment analysis was performed as described in Horta et al^4^.

For pairwise analysis of Greek Island interactions (Figures 3c and 3d) the HiC contacts mapping to the 50 Kb bin containing each Greek Island were included. One Greek Island, Amorgos, spans the border of two 50 Kb bins; both were included (“Amorgos A” and “Amorgos B”). Boxplot in figure 3d displays the range of values observed for trans contacts made by each Greek Island; whiskers indicate the 1.5 * the interquartile range and outliers are not shown. In order to quantify the effect of Ldb1KO on OR clusters and Greek Islands (Figure 3f), all 50 Kb bins overlapping each OR cluster were identified. These OR cluster bins were subdivided based upon whether or not they were associated with a Greek Island. Greek Islands exhibit increased contacts over a broad area (see HiC contact heatmaps in Figure 3a and 3b), so bins were assigned to the Greek Island group if they overlapped or were within 50kb of a Greek Island. In total, 768 bins overlapping OR clusters were identified, out of which 136 were assigned to the Greek Island set and 632 were assigned to the OR cluster set. The HiC score was determined for every *trans* pair of bins (176,079 OR cluster to OR cluster pairs, 8,413 Greek Island to Greek Island pairs). In addition, a set of random genomic regions was generated for comparison.

This random set was generated by shuffling the genomic location of OR clusters. Each OR cluster was moved to a random new position on the same chromosome. The randomly shuffled OR clusters were checked to ensure they did not overlap any existing OR clusters. 50 Kb bins overlapping the shuffled OR clusters were then identified and analyzed as described above. The cumulative distribution of HiC scores for each of these three groups was then plotted for Control OSNs and Ldb1KO OSNs. For plots of average Greek Island HiC contacts (extended data 4c), all of the HiC scores for contacts made by each Greek Island with *trans* Greek Islands were averaged, and these data were fit with a linear model with the y-intercept set to 0. A similar approach was used for OR clusters and or cluster trans contacts (extended data 4d), but in addition all bins comprising each cluster (excluding Greek Island overlapping and adjacent bins) were averaged together to generate an average score for each cluster.

Juicer Tools was used to generate APA plots and calculate APA scores for *trans* interactions between Greek Islands. APA analysis was run at 10kb resolution with 20 flanking windows. The normalized APA matrix was plotted with the scale set to 5 times the mean of the matrix.

In order to generate signal tracks for all *trans* Greek Island contacts (extended data 5a,b) the HiC signal anchored on each Greek Island located in trans was added together to generate a single plot for each chromosome. Signal plots were visualized using the UCSC Genome Browser. For extended data 6b, HiC signal tracks anchored on the H enhancer were generated with Juicebox and then plotted with Deeptools2. Values are KR normalized counts per billion HiC contacts. Each row of the heatmap is a 2 Mb region centered on a Greek Island located in trans relative to H. Some intervals overlap due to the presence of multiple Greek Islands within 1 Mb of each other. Signal plots present average data for all peaks. Heatmaps are sorted by mean signal in Control OSNs.

### RNA-seq

Control OSN RNA-seq data, prepared from GFP+ cells purified from *OMP-IRES-GFP* mice, were previously published^5^ (GSE93570). Ldb1KO OSNs were FACS-purified based upon expression of Cre inducible tdtomato+, and RNA-seq libraries were prepared from FACS-purified live cells as previously described. See RNA-seq tab of Supporting Information 1 for a summary of RNA-seq sequencing data.

CutAdapt was used to remove Adapter sequences from raw sequencing data and then filtered reads were aligned to the mouse genome (mm10) using STAR^41^ v2.5.3a. Samtools was used to select uniquely aligning reads by removing reads with alignment quality alignments below 30 (-q 30). RSeQC^42^ (RRID:SCR_005275) was used to generate RNA-seq signal tracks with signal normalized to a library size of 10,000,000 reads.

RNA-seq data analysis was performed in R with the DESeq2^43^ package. Very low abundance transcripts (genes with fewer than 10 counts combined across all samples) were excluded. DESeq2 was used to calculate FPKM values for Control OSNs and Log2 fold change values for comparisons of Control and Ldb1KO OSNs.

### Immunofluorescence

MOE was dissected from 6-week Ldb1KO (Ldb1fl/fl;OMPcre) mice and littermate controls. MOE tissue was embedded in OCT and then coronal cryosections were collected at a thickness 12uM. Tissue sections were prepared and stained as previously described^5^. Tissue sections were stained with primary antibodies for Ldb1 (1:1000 dilution, Santa Cruz Biotechnology Cat# sc-11198, RRID:AB_2137017), Adcy3 (1:200 dilution, Santa Cruz Biotechnology Cat# sc-588, RRID:AB_630839), and Olfr1507 (1:2000 dilution, courtesy of Gilad Barnea). DNA was labeled with DAPI (2.5ug/mL, Thermo Fisher Scientific Cat# D3571). Primary antibodies were labeled with the following secondary antibodies: for Ldb1, anti-goat IgG conjugated to Alexa-488 (2ug/mL, Thermo Fisher Scientific Cat# A-11055, RRID:AB_2534102), for Adcy3, anti-rabbit IgG conjugated to Alexa-555 (2ug/mL, Thermo Fisher Scientific Cat# A-31572, RRID:AB_162543), for Olfr1507, anti-guinea pig IgG conjugated to Cy3 (2ug/mL, Jackson ImmunoResearch Labs Cat# 706-165-148, RRID:AB_2340460). Confocal images were collected with a Zeiss LSM 700 and image processing was carried out with ImageJ (NIH).

### Statistics

For ChIP-seq, statistically significant peaks were identified using HOMER on each replicate of each experiment. Candidate peaks were selected by setting a read count threshold based upon an input control false discovery rate of 0.001, and then peaks were filtered based upon the following criteria: Poisson p-value over input < 1.00e-04 and Poisson p-value over local region < 1.00e-04. Consensus peak sets were then generated by selecting peaks that overlapped in at least two biological replicates. For HiC data, two independent replicates were generated for each condition and analyzed separately. Individual biological replicates yielded similar results and were pooled for the analyses presented here. For RNA-seq, three biological replicates of Control OSNs and two biological replicates of Ldb1KO OSNs were analyzed with DESeq2, which generates two-tailed Wald test p-values, and generates adjusted p-values using the Benjamini-Hochberg method.

### Data Availability

The ChIP-seq and RNA-seq data reported in this paper are available from GEO (GSE112153). HiC data are available from the 4D Nucleome project (4DNESEPDL6KY, 4DNES425UDGS, 4DNESRE7AK5U, 4DNES54YB6TQ, 4DNESNYBDSLY, 4DNESH4UTRNL). Control OSN RNA-seq, Lhx2 ChIP-seq, and Ebf ChIP-seq data were previously described^5^ and are available from GEO (GSE93570).

---

**Extended Data Figure 1:**
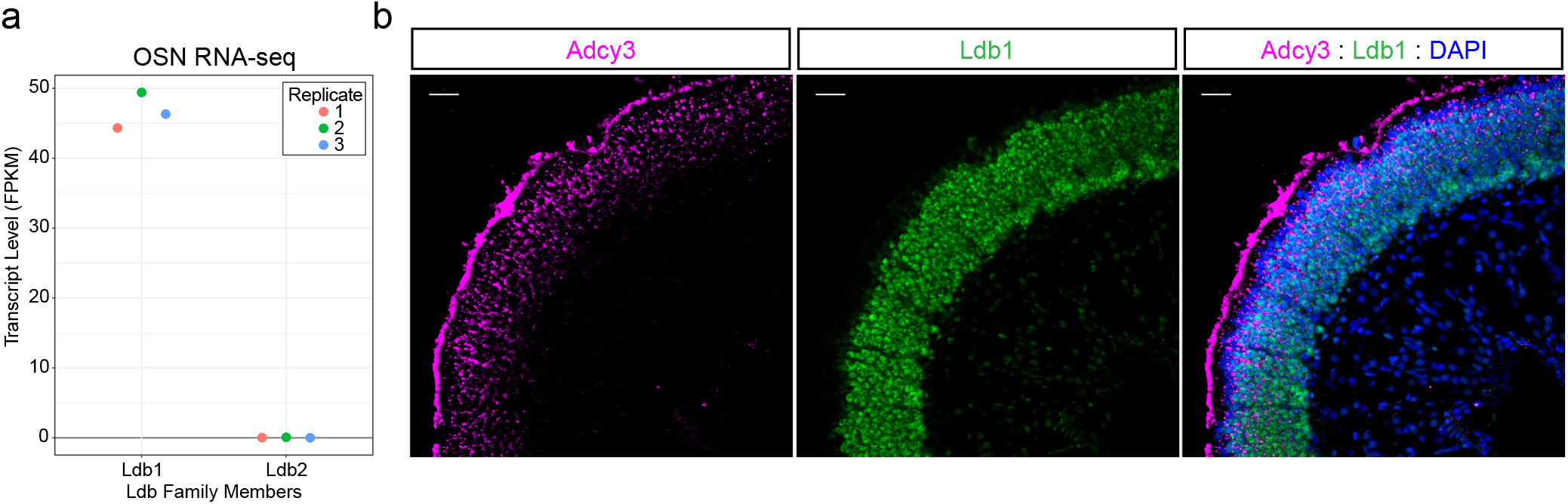
Expression of Ldb family members in OSNs. **a)** Transcript level of both Ldb family members in 3 independent Control OSN RNA-seq data sets. **b)** Sections of olfactory epithelium stained for Ldb1 (green) and Adcy3 (magenta), a marker for mature OSNs. Nuclei are labeled with DAPI (blue). Scale bar = 25um.

**Extended Data Figure 2:**
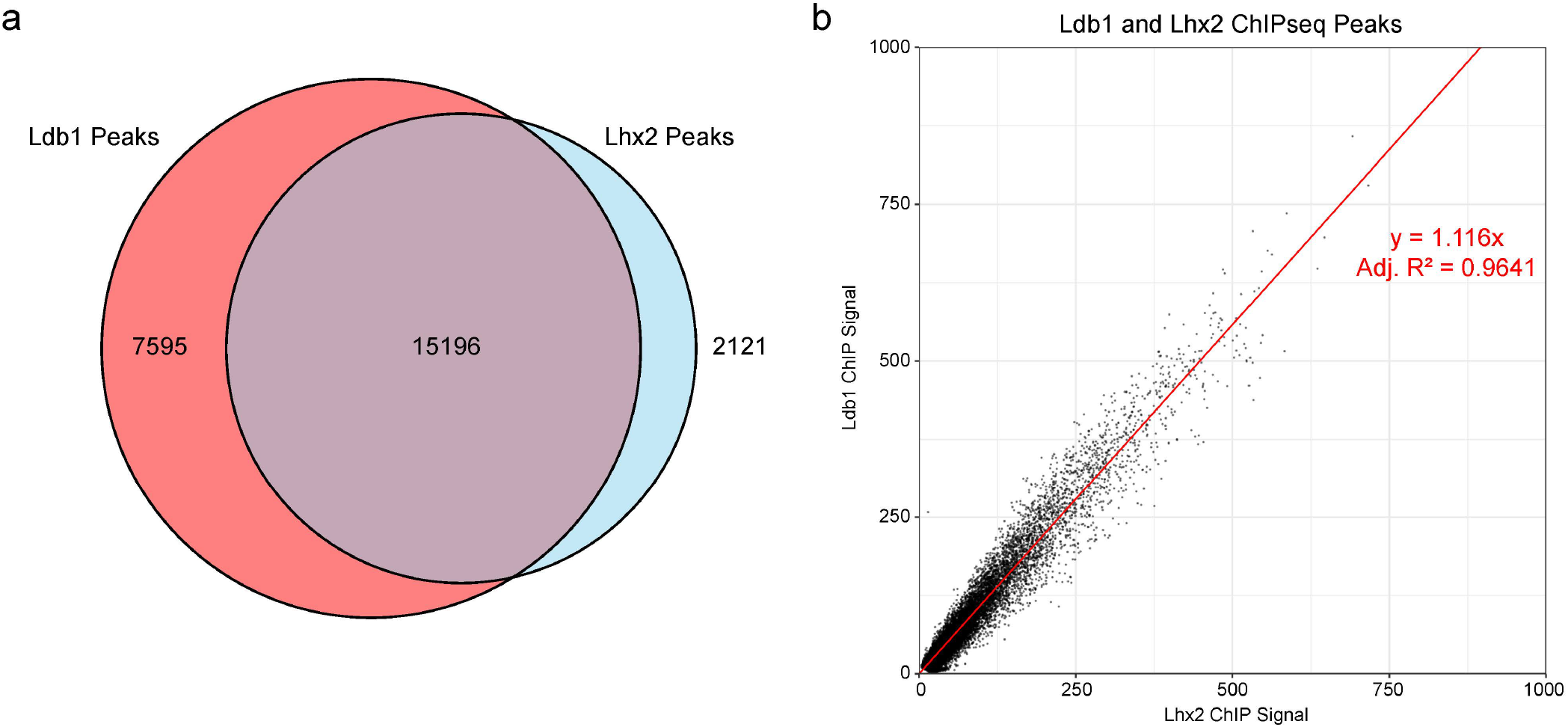
Highly correlated binding of Lhx2 and Ldb1 in OSNs. **a)** Extensive overlap between consensus Lhx2 and Ldb1 ChIP-seq peak sets. **b)** Very close relationship between normalized Lhx2 ChIP signal and Ldb1 ChIP signal. All peaks observed with either Lhx2 or Ldb1 are plotted together with a best fit line obtained by linear regression with y-intercept set to 0.

**Extended Data Figure 3:**
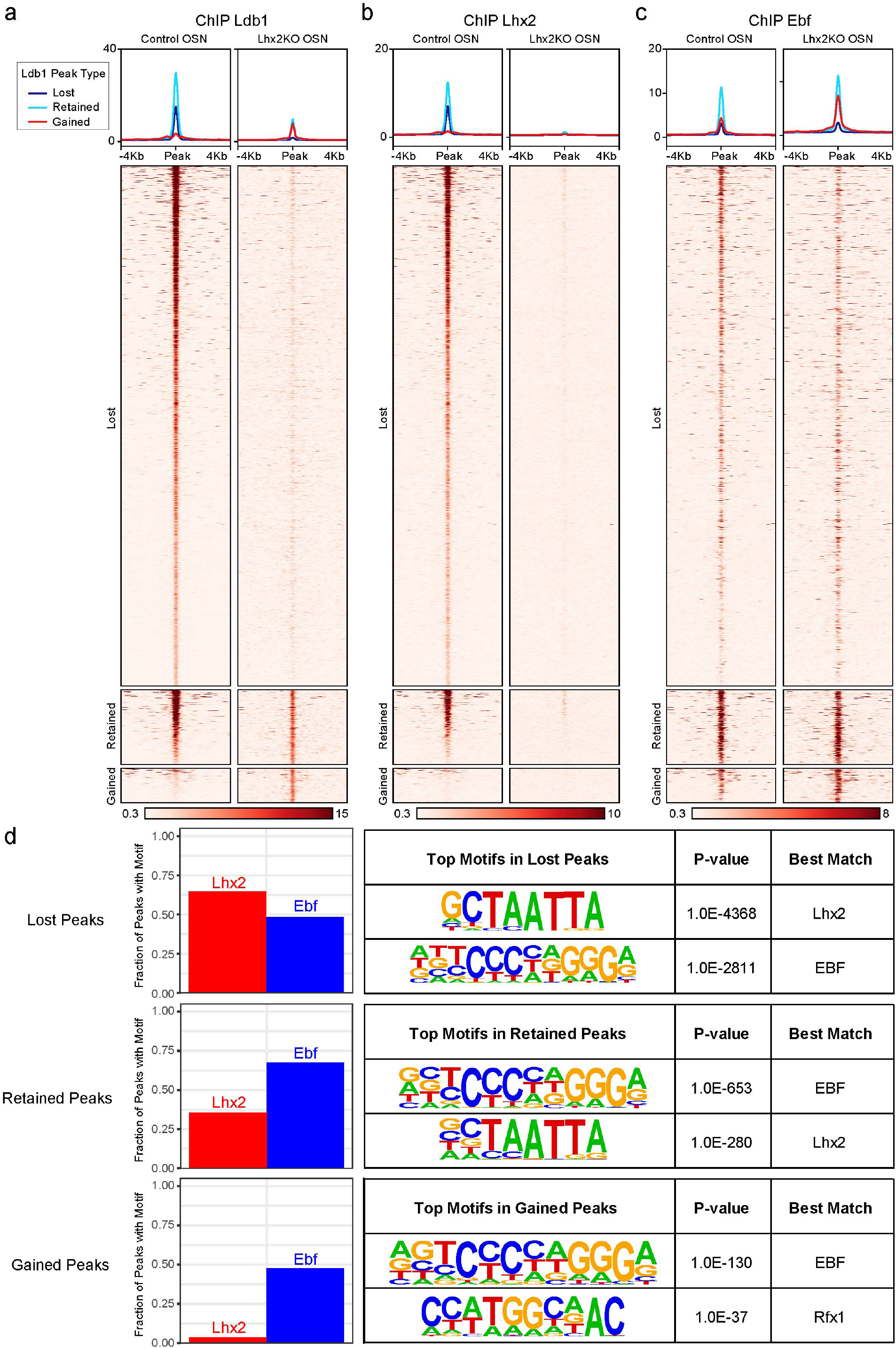
Loss of Lhx2 uncovers co-binding of Ldb1 and Ebf. **a)** Ldb1 ChIP signal in control versus Lhx2 KO OSNs. Plot shows average signal across all sites and heatmap shows individual peaks. Peaks are grouped by whether they are observed only in control OSNs (“Lost” in Lhx2KO), in both populations (“Retained” in Lhx2KO), or only in Lhx2KO OSNs (“Gained” in Lhx2KO). **b)** Lhx2 ChIP signal from control OSNs and Lhx2KO OSNs is shown for Ldb1 peaks from **a**. **c)** Ebf ChIP signal from control OSNs and Lhx2KO OSNs is shown for Ldb1 peaks from **a**. 773 out 1324 (58.4%) of “Gained” Ldb1 peaks overlap an Ebf ChIP peak observed in Lhx2KO OSNs. **d)** Motif analysis of peaks in the “Lost”, “Retained”, and “Gained” categories. Left: Fraction of peaks containing an Lhx2 or Ebf motif. Right: Top two de novo motifs with p-value and identity of the most similar known motif.

**Extended Data Figure 4:**
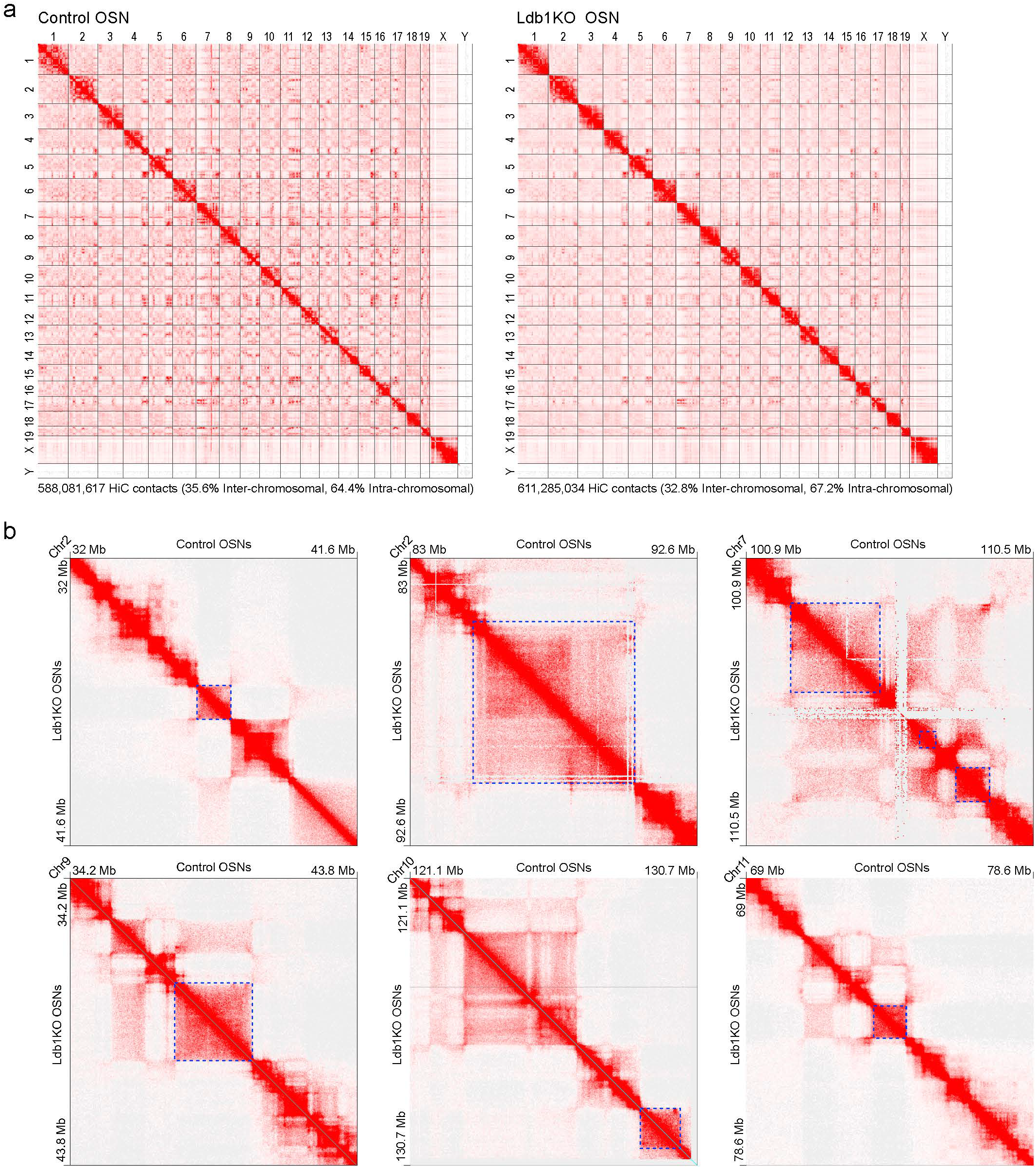
Overall nuclear architecture is unchanged in Ldb1KO OSNs. **a)** Genomewide HiC profiles of Control and Ldb1KO OSNs. b) HiC contacts in Control and Ldb1KO OSNs (25kb resolution) for 6 regions containing OR Clusters (blue boxes).

**Extended Data Figure 5:**
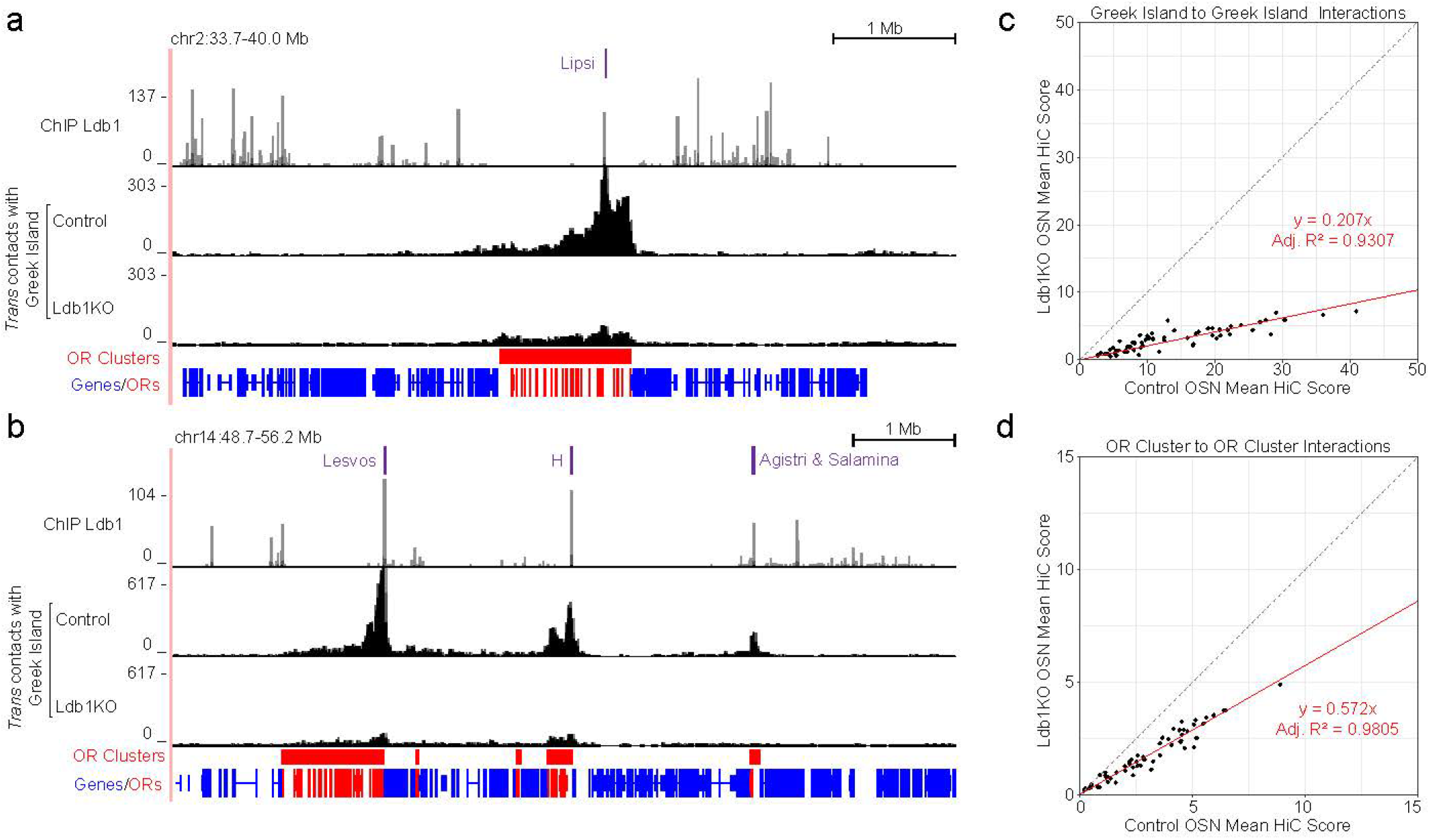
Loss of Ldb1 eliminates focal contacts between Greek Islands. **a+b)** Signal plot shows HiC contacts made by *trans* Greek Islands across multiple OR clusters (25kb resolution). In Ldb1KO cells contacts with trans Greek Islands are greatly reduced, particularly at site of Greek Island to Greek Island contacts. **c)** For each Greek Island, the mean HiC score for *trans* contacts with other Greek Islands is plotted for Control OSNs and Ldb1KO OSNs (50kb resolution). **d)** As in c, except for mean HiC score is shown for each OR Clusters contacting *trans* OR clusters.

**Extended Data Figure 6:**
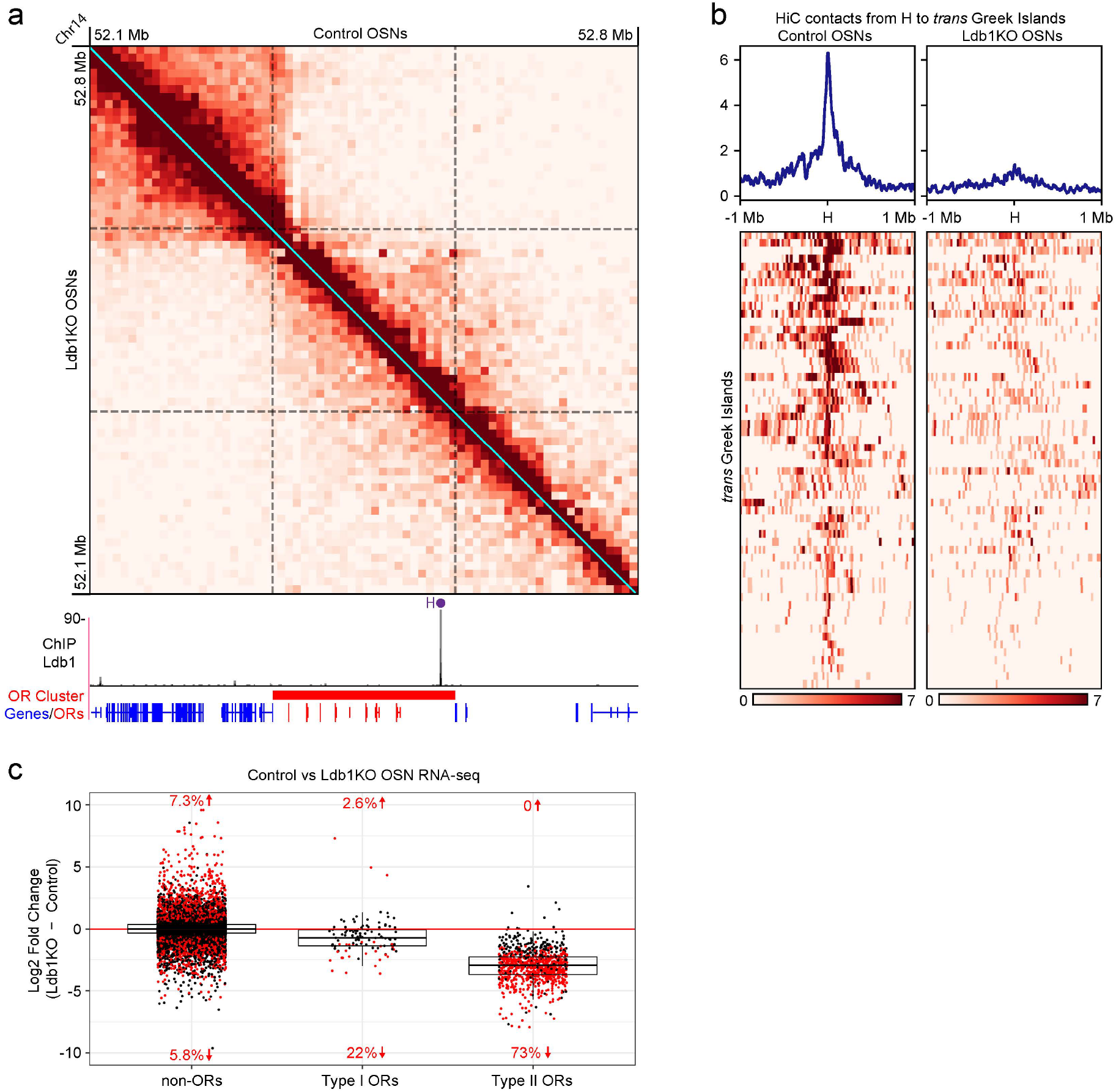
Loss of contacts between H and Greek Islands in Ldb1KO OSNs. **a)** HiC contacts (10kb resolution) surrounding the Olfr1507 locus and H Enhancer are similar in Control and Ldb1KO OSNs. **b)** Loss of Ldb1 results in reduced interactions between H and *trans* Greek Islands. Plot shows average contacts between the 25kb bin containing H and a 1MB region centered on each *trans* Greek Island, and the heatmap shows each individual interaction. **c)** Ldb1KO reduces transcript levels of Type II ORs much more than Type I ORs.

